# Survival, Movement, and Lifespan: Decoding the Roles of *Patched-Related* (*Ptr*) in *Drosophila melanogaster*

**DOI:** 10.1101/2024.05.07.592982

**Authors:** Cristina Parada, Daniel Prieto

**Affiliations:** Departamento de Neurofarmacología Experimental, Instituto de Investigaciones Biológicas Clemente Estable, Montevideo, Uruguay; Programa de Desarrollo de las Ciencias Básicas (PEDECIBA), Uruguay; Departamento de Neurofisiología Celular y Molecular, Instituto de Investigaciones Biológicas Clemente Estable, Montevideo, Uruguay; Departamento de Biología del Neurodesarrollo, Instituto de Investigaciones Biológicas Clemente Estable, Montevideo, Uruguay

**Keywords:** Patched-related (Ptr), development, survival, muscle, larval locomotion, lifespan, Drosophila

## Abstract

Patched-related (Ptr) is a transmembrane protein implicated in developmental processes in *Drosophila melanogaster*, yet its precise role remains incompletely understood. Here, we use *Ptr*^*23c*^ null mutants to investigate the functional significance of Ptr through the entire life cycle monitoring survival during embryonic, larval, pupal and adult development, and studying larval locomotion and muscle structure. We report that *Ptr*^*23c*^ larvae displayed impaired hatching, indicative of defective embryonic development. Moreover, mutant larvae exhibited reduced mobility and lethargy, suggesting a potential involvement of Ptr in neuromuscular function. Morphological analysis of somatic muscles in mutant larvae revealed enlarged cell nuclei. Despite high pre-adult mortality, a subset of *Ptr*^*23c*^ mutant adults display an unexpected extension in lifespan compared to controls, implicating Ptr in the regulation of longevity. Our findings provide critical insights into the multifaceted role of Ptr in *Drosophila* development, highlighting its contributions to post-embryonic survival, neuromuscular function, and lifespan regulation. This study underscores the significance of exploring broader genetic networks to unravel the complexities of developmental processes.

**Highlights:**

- Loss of Ptr elicits impaired embryonic hatching, reduced mobility, and lethargy.
- Ptr23c mutants display nuclear enlargement in somatic muscles.
- Ptr23c individuals that survive to adulthood demonstrate extended longevity.

## Introduction

Patched-related (Ptr) is a multipass transmembrane protein related to the canonical receptor of the Hedgehog (Hh) pathway, Patched (Ptc) (Pastenes et al., 2008) encoded by the evolutionarily conserved gene *Ptr*, which is ubiquitously expressed at every stage of *Drosophila melanogaster* development (Bolatto et al., 2022, 2015; Pastenes et al., 2008). The expression of *Ptr* begins at the cellular blastoderm (Pastenes et al., 2008; Zúñiga et al., 2009) and reaches its highest level later during embryonic development (modENCODE, Brown et al., 2014). During gastrulation, the expression of *Ptr* mRNA follows the typical pattern of pair-rule genes (Zúñiga et al., 2009) and at later stages is expressed in several tissues, including hemocytes (Bolatto et al., 2015; Cattenoz et al., 2020).

Results from an *in vitro* assay suggested that Ptr could compete with Ptc for the ligand Hh, serving as an additional modulator of Hh signaling (Bolatto et al., 2022). While PTCHD-18, a *C. elegans* ortholog interacts with the Hedgehog-related protein promoting its removal from the extracellular milieu (Chiyoda et al., 2021), the human ortholog PTCHD1 binds cholesterol but not Sonic Hedgehog (Hiltunen, et al., 2023). Absence of Ptr either in the *Ptr*^*23c*^ mutant or after silencing the gene causes high embryonic mortality and nervous system alterations in *D. melanogaster* (Bolatto et al., 2022; Zhao et al., 2008), but the precise molecular function of Ptr remains elusive. Our investigation sought to elucidate specific contributions of *Ptr* to *Drosophila* development, with a particular emphasis on its effects on post-embryonic survival, larval locomotion, and adult lifespan.

We employed the null mutant *Ptr*^*23c*^ (Bolatto et al., 2015), and found that most embryos fail to hatch. Among the few individuals that survive to the larval stage, we observed lethargy and abnormal muscle nuclei. Despite these challenges, a small proportion of *P tr*^*23c*^ mutants manage to develop to the adult stage and notably, they exhibit extended lifespan. Our findings shed light on the intricate interplay between *Ptr*function and the phenotypic outcomes observed throughout the life stages of *Drosophila*.

## Materials and Methods

### Genetics and Drosophila strains

*Drosophila melanogaster* sotcks were obtained from the following sources: *w; Df(2L)32FP-5/CyO*,*twist-Gal4*,*UAS-GFP* (Pirone 2016); *w*; He-GAL4*.*Z, UAS-GFP*.*nls* (#8700, BDSC); *w*, Bl/CyO; TM2/TM6b* balancer, *Ptr*^*23c*^*/CyO, ftz-lacZ* (Bolatto et al., 2015). Standard crosses were performed to generate the *Ptr*^*23c*^*/CyO, twi-GFP* and the *Ptr*^*23c*^*/CyO, twi-GFP; He-GFP/He-GFP* stocks. Wild-type Oregon-R controls were used unless otherwise stated.

Flies were maintained on standard medium, as previously described (Silvera et al., 2025). Larvae selected for experiments were cultured in apple juice-agar medium supplemented with yeast.

### Embryo collection

Eggs layed on apple juice-agar plates supplemented with yeast were delicately collected using a brush and subsequently washed with tap water. De-chorionation was achieved by immersion in 5% sodium hypochlorite, followed by rinsing with PBST (0.05% Triton X-100 in PBS). Embryonic stages were determined according to morphological criteria (Campos-Ortega and Hartestein, 1997). When necessary, embryos were selected based on GFP fluorescence (see below).

### Hatching index

Groups of 30 late stage 17 embryos were transferred to apple juice-agar plates. The number of larvae was counted after 24 hours and the hatching index was determined as Hatching index = Hatched larvae/Total larvae.

### Measurement of larval locomotion

Larvae were recorded during 45 sec on video under the dissecting microscope. Crawled distance and crawling velocity were assessed using the Manual Tracking plugin from FIJI, following methods outlined in Balakrishnan et al., 2021.

### Measurement of lifespan

Experiments were performed with *+/+; +/+; He-GFP/He-GFP* control, *+/+; Ptr*^*23c*^*/Ptr*^*23c*^; *He-GFP/He-GFP* mutant and *+/+; Ptr*^*23c*^*/+; He-GFP/He-GFP* heterozygous individuals. For measurements, 128 males of the control strain were kept in 12 vials; 22 *Ptr*^*23c*^ mutant males were kept in 4 vials and 118 males of the heterozygous strain were kept in 12 vials. The dead flies were counted every 2 days, and thereafter the living flies were transferred to fresh food every 2–4 days.

### Staining of larval somatic muscles

Third-instar larvae were dissected along the dorsal midline in ice-cold PBS. The viscera were removed, and body-wall preparations were stretched and incubated in 0.5 mM EGTA for 5 min. Body walls were fixed in 4% paraformaldehyde (PFA) at room temperature for 15 minutes, washed twice with PBS-T (0.3% Triton X-100 in PBS), incubated overnight at 4°C with DAPI (1 μg/ml) and eosin-conjugated Phalloidin (1:200, Molecular Probes), and mounted in 80% glycerol (in 1 mM Tris-HCl, pH=8). Images of VL3 muscles of the 5^th^ abdominal segment were taken for nuclear area quantification.

### Microscopy and Imaging

Fluorescent embryos were selected using a Wesco m300 series stereo microscope (USA) with a Nightsea SFA fluorescence adapter (Nightsea, Hatfield, PA, USA). Locomotion was imaged using a mobile phone camera with a 3D-printed ocular adapter (https://www.instructables.com/Universal-Phone-MountAdapter-for-Microscope-Telesc/).

Muscles were imaged with a Zeiss LSM800 laser-scanning confocal microscope using an EC Plan-Neofluar 20x/0.50 M27 objective and a 1.3X zoom using a GaAsP-PMT detector. Z-stacks of 1024 x 1024 px images were taken at 500 nm steps. Subsequent image processing was performed using FIJI. Nuclear area was quantified on maximum intensity projections of z-stacks. Figures were crafted using Inkscape 1.1 (Inkscape Project, 2020) and GIMP 2.10 (The GIMP Development Team, 2019).

### Quantification and Statistical Analysis

Data analyses were conducted using GraphPad Prism version 9.5. Mann-Whitney U-tests with a significance level set at p<0.05 were performed. Survival probabilities were calculated using Gehan-Breslow-Wilcoxon and Mantel-Cox tests.

## Results

### Most *Ptr*^*23c*^ mutants die when trying to hatch

We began focusing on the effect of the null mutation *Ptr*^*23c*^ on survival through multiple developmental stages. Available data indicates that *Ptr* is essential for embryonic survival. *Ptr*^*23c*^ mutants generated by P-element excision (Bolatto et al., 2015) exhibit a mortality rate of 81% during embryogenesis (Bolatto et al., 2022). We found that RNAi-induced knockdown of *Ptr* under the control of a maternally expressed alphaTub67C promoter caused between 10% to 46% of mortality during embryonic development (data not shown). Later in embryogenesis, by stage 17, *Ptr*^*23c*^ embryos exhibited a reduced hatching index compared to control embryos of the same age (median: control 96%, mutant 79%, p < 0.05) (Fig. 1A). Careful examination revealed that mutants lived for up to 24 additional hours and showed the peristaltic contractions characteristic of pre-hatching, but failed to hatch and died within the egg shell.

**Figure 1.**
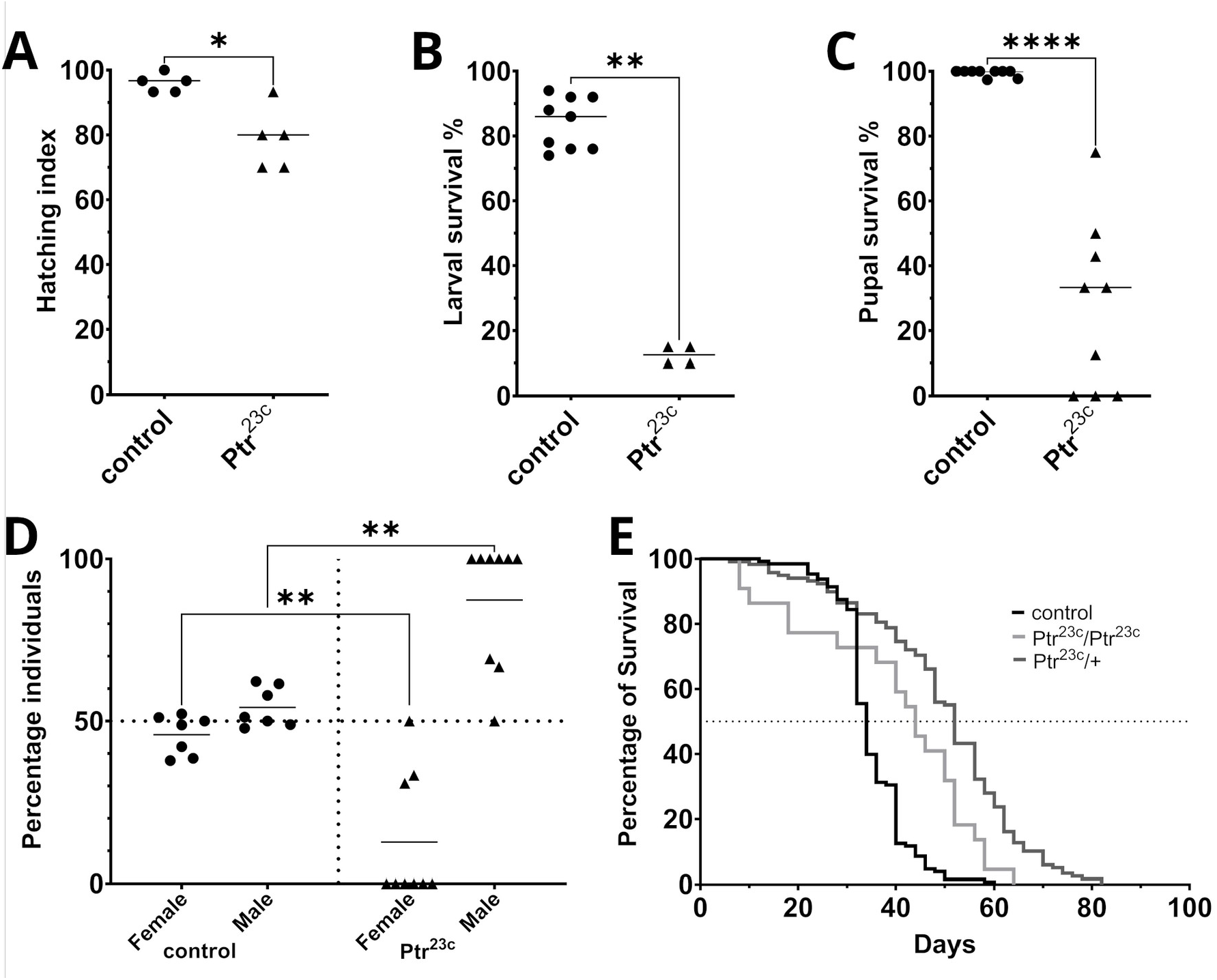
The *Ptr*^*23c*^ mutation affects survival at every stage of the life cycle of *Drosophila*. **(A)** Reduced larval hatching among *Ptr*^*23c*^ mutants. Hatching index was calculated and the median is compared: control 96%, mutant 79%; n = 5 plates with 30 embryos each; p < 0.05. **(B)** Reduced survival of larvae among *Ptr*^*23c*^ mutants. A very low proportion of mutant larvae reach the pupal stage. Median: control 83%, mutant 13%; n = minimum of 4 plates with 40 larvae each; p < 0.01. **(C)** Reduced mutant survival along the pupal stage. Median: control 100%, mutant 33%; p < 0.0001, assessed by Mann-Whitney test. n = 9 cages with pupae each. **(D)** At the pupal stage the proportion of individuals of each sex showed a significant sex disparity in the mutant. Mean: control males 54%, females 46%; mutant males 87%, females 13%; n = 7 (control) and 9 (mutant) cages each. p < 0.01. **(E)** *Ptr*^*23c*^ mutant adult males exhibited extended lifespan (median: 50 days) compared to control males (median: 34 days). Significant differences revealed by Kaplan-Meier analysis (p < 0.0001) in Gehan-Breslow-Wilcoxon and Mantel-Cox tests; Chi-squared = 59.6. N = 100 *+/+; +/+; He-GFP/He-GFP* (control), 22 *+/+; Ptr*^*23c*^*/Ptr*^*23c*^; *He-GFP/He-GFP* (Ptr^23c^/Ptr^23c^), and 118 *+/+; Ptr*^*23c*^*/+; He-GFP/He-GFP* (Ptr^23c^/+).

### Lower survival in *Ptr*^*23c*^ larvae and pupae contrast with longer adult lifespan

When raised in standard culture medium, most *Ptr*^*23c*^ larvae die before molting into the second instar. We found that it was possible to keep these mutants alive by culturing them on a soft apple juice-agar medium supplemented with yeast. Although mutant larvae experienced reduced survival, some individuals completed larval development and reached the pupal stage (median: control 83%, mutant 13%, p < 0.01) (Fig. 1B). A significant decrease in survival was also observed at the pupal stage when mutants were compared with control pupae (median: control 100%, mutant 33%, p < 0.0001), as determined by the Mann-Whitney test (Fig. 1C). Some mutant pupae reached the final stage of pre-adult development as fully formed pharate adults.

Remarkably, a sex effect of the *Ptr*^*23c*^ mutation was observed, with a much lower proportion of females reaching this stage (mean: control males 54.2% and females 45.8%; mutant males 87.3%, females 12.7%, p < 0.01) (Fig. 1D). The exceptionally low proportion of mutant females reaching the adult stage posed a challenge for assessing lifespan in individuals of this sex. Consequently, our analysis of adult lifespan was limited to males.

The small subset of *Ptr*^*23c*^ mutants that survived into adulthood exhibited an extended lifespan compared with control flies (median = 44 days vs. 34 days). Heterozygous *Ptr*^*23c*^/+ individuals had longer survival time (median = 52 days). Kaplan-Meier analysis revealed significant differences (p < 0.0001) in both the Gehan-Breslow-Wilcoxon and Mantel-Cox tests (Fig. 1E).

### Locomotion is impaired at larval stages

We observed that *Ptr*^*23c*^ larvae exhibited a lethargic behavior. When assayed with a locomotion test early in larval development (first instar) mutant larvae had a significantly reduced locomotion (19.8 mm) compared to control larvae (36.5 mm, Fig 2A). Additionally, mutant larvae moved at a 54% slower median speed (0.45 mm/sec) compared with control larvae (0.83 mm/sec Fig. 2B). Towards the end of larval life (third instar), mutant larvae continued to show deficient locomotion (154.5mm; 3.51 mm/sec) relative to control larvae (224.8; 5.11 mm/sec), although the differences were diminished to a 68% from that of the control.

**Fig 2:**
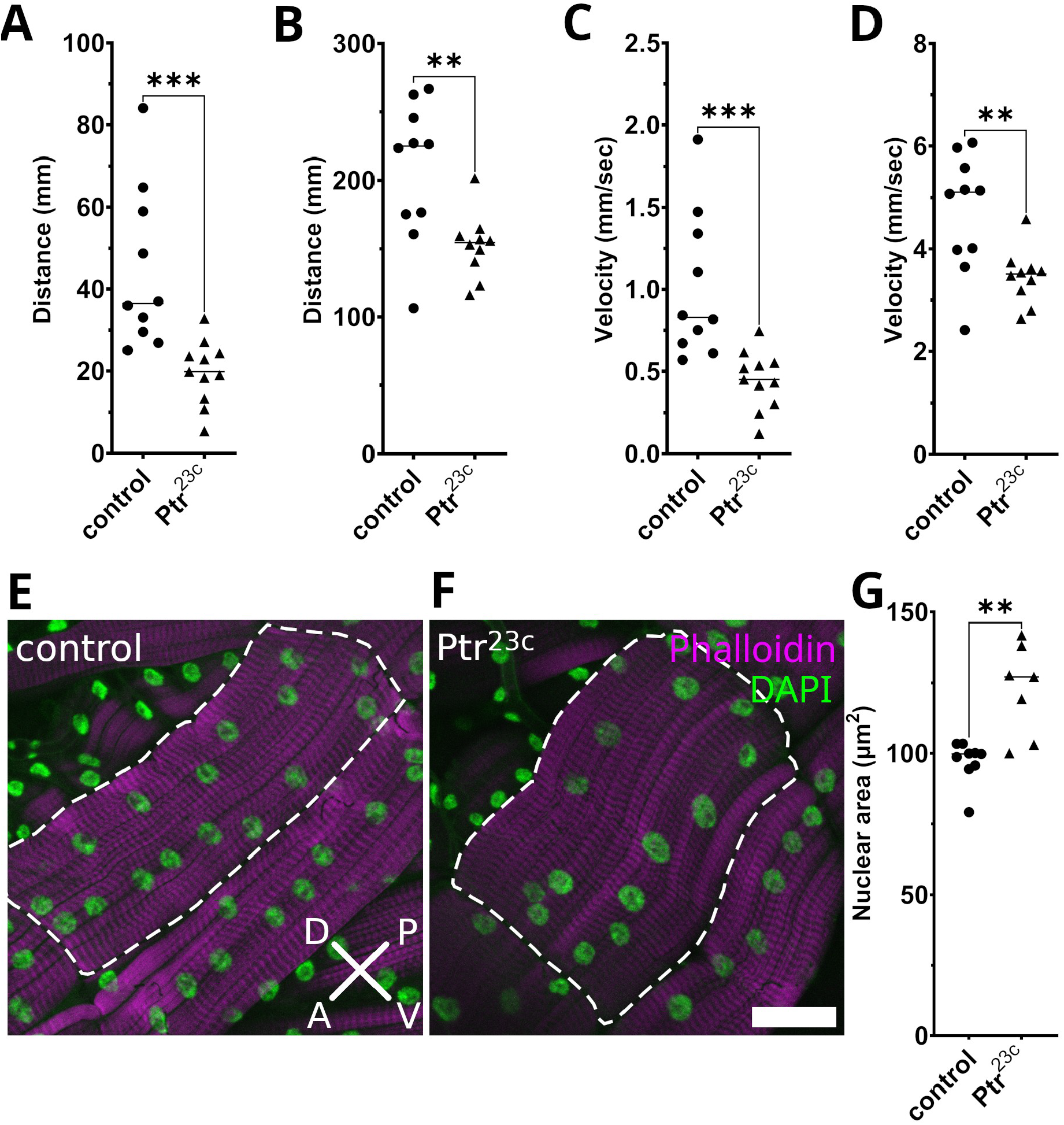
Locomotion and muscular phenotype associated with the *Ptr*^*23c*^ mutation. Motor impairment in *Ptr*^*23c*^ mutant larvae **(A-D)** correlates with a muscular phenotype **(E -G). (A)** The *Ptr*^*23c*^ mutation reduced the crawling distance covered by first-instar larvae during 45 sec from 27.4 mm in control larvae (n=10) to 19.8 mm in *Ptr*^*23c*^ (n=11). **(B)** Third-instar mutant larvae showed a reduction of crawling distance from 224 mm, 160.5 mm, n=10 (control larvae) to 154 mm, 85.6 mm, n=10. Crawling velocity was also reduced **(C)** from 0.83 mm/sec (control larvae) to 0.45 mm/sec in first instar larvae and **(D)** from 5.11 mm/sec (control larvae) to 3.51 mm/sec in third-instar larvae (Medians are shown in A to D). Statistical significance was determined using the Mann-Whitney U-test, yielding a p-value < 0.001 and 0.01. **(E-F)** Confocal laser scanning microscopy images of somatic muscle VL3 (dashed line) showing nuclei in green and F-actin in magenta. Maximum intensity projection images from a representative control larva **(E)** and a *Ptr*^*23c*^ mutant larva (**F**). Scale bar: 50 µm (20 planes at 0.5 μm intervals along the z axis). **(G)** Nuclear size evaluation through the measurement of nuclear area on maximum intensity z-projections of confocal stacks from body wall preparations. Statistical analysis, performed using the Mann-Whitney U-test, revealed that the area of muscle nuclei was 27% larger in the mutant compared to control. Median: control 99.8 μm^2^, mutant 127.1 μm^2^; p-value < 0.01. VL3 abdominal muscle was used for quantification. n= 9 control and 7 mutant larvae. IQR: Interquartile range; A: anterior; P: posterior; V: ventral; D: dorsal.

### Mutant larvae exhibit enhanced endoreplication of somatic muscle nuclei

Hatching and locomotion depend on the interplay between the nervous system and musculature (Baker et al., 2018; Crisp et al., 2008; Hunter et al., 2021). While previous studies have reported abnormal nervous system morphology in *Ptr* mutants (Zhao et al., 2008; Bolatto et al., 2022), the potential impact on musculature has not been explored. Given that *Ptr* is known to be expressed in muscle cells (Xiao et al., 2023), we employed F-actin and DNA staining to examine the somatic muscles of *Ptr*^*23c*^ third-instar larvae. Our analysis revealed that the mutation caused enlargement of cell nuclei across all muscles (Fig. 2E-G, median area: control = 99.8 μm^2^, *Ptr*^*23c*^ = 127.1 μm^2^).

## Discussion

The critical role of *Ptr* during early development in *Drosophila melanogaster* is underscored by the high mortality rate observed in *Ptr*^*23c*^ mutants during embryogenesis (Bolatto et al., 2022). To investigate the effects of the *Ptr*^*23c*^ mutation across the entire life cycle, we developed a culture strategy that enabled some mutant individuals to survive larval and pupal development and reach adulthood. Through a comprehensive analysis of the post-embryonic mutant phenotypes we propose that *Ptr* interacts with the insulin pathway.

We observed clear impairments in both hatching from the eggshell and locomotion during larval development. These biological processes depend on well-coordinated, strong muscle contractions controlled by the nervous system through sophisticated motor programs (Crisp et al., 2008; Baker et al., 2018; Hunter et al., 2021). Both phenotypes point to a neuromuscular dysfunction and offer a plausible explanation for the high mortality observerd in *Ptr*^*23c*^ mutants during embryonic and larval stages. An association between abnormal nuclear size and muscle function has been reported in *Drosophila* larvae with mutations in other genes (Wang et al., 2018), making it likely that the observed correlation between enlarged myonuclei and impaired muscle function in *Ptr*^*23c*^ mutants reflects a functional interaction. Moreover, our finding of abnormal muscle morphology aligns with previous reports of abnormal nervous system morphology in *Ptr*^*23c*^ mutants (Zhao et al., 2008; Bolatto et al., 2022), strengthening the case for the possibility that *Ptr* absence affects the structure and function of both the nervous and muscular systems.

While not addressed in this study, we hypothesize that the increased mortality of mutant pupae may result from impaired neuromuscular function, as ecdysis requires coordinated muscle contractions to break open the pupal case (Kim et al., 2006; Elliott et al., 2021). Interestingly, we observed mortality at both early and late pupal stages (data not shown), with the highest mortality detected in *Ptr*^*23c*^ female pupae.

Although we did not explore causation, the sex-specific differences in adult mortality appear to correlate with differences in *Ptr* expression. Specifically, higher *Ptr* expression has been reported in the central nervous system of females, as shown by RNAseq analysis (Leader et al., 2018, *flyatlas2*.*org*).

Sexual dimorphism in *Drosophila* is regulated by various genes. One key factor, the female sex determinant *Sex-lethal*, is expressed in two distinct neuronal subpopulations within the central nervous system, one of which corresponds to insulin-producing cells. This sex-specific regulation of insulin-like peptide production may influence tissue growth globally (Sawala and Gould, 2017). Additionally, insulin-like peptides are known to play a crucial role in the development of the pupal nervous system and ecdysis (Truman and Riddiford, 2023).

The contrast between high pre-adult mortality and extended adult lifespan among the few escapers in *Ptr*^*23c*^ mutants prompts further exploration of the complex interplay of factors shaping their life history. The increased survival observed in heterozygotes compared to homozygotes may reflect to the suppression of deleterious effects during earlier developmental stages. Alternatively, it could result from more intricate interactions with other components of the cellular pathways in which Ptr is involved, as has been reported for other genes (Bai et al., 2015). Genome-wide analysis of single-nucleotide polymorphisms has identified *Ptr* in one of the genomic regions associated with longevity (Durham et al., 2014).

Nutrient signaling pathways are known to influence lifespan in *Drosophila*, with signals secreted from the gut, fat and muscle tissues triggering protective effects across the organism (reviewed in Piper and Partridge, 2018). On the other hand, Ptr has been proposed as a potential repressor of the Hh pathway (Bolatto et al., 2022).

Since increased expression of Hh extends lifespan (Rallis et al., 2020), the longer lifespan in *Ptr*^*23c*^ mutants may be linked to the activation of this pathway. Our unexpected observation of extended adult life within *Ptr* mutant males raises intriguing questions about the underlying mechanisms and potential connections with the metabolic processes commented below.

The muscular phenotype observed in *Ptr*^*23c*^ mutants suggests a potential link between Ptr-mediated Hh signaling and muscle development. During larval growth, somatic muscles expand rapidly, accompanied by increased nuclear endoreplication, which is tightly regulated by a complex interplay of the Hh, insulin receptor (InR) and dFOXO pathways (Demontis & Perrimon 2009; Singh et al., 2018). In ovary follicle stem cells, antagonistic effects between the insulin and Hh signaling pathways have been observed, with Hh dependent autophagy being inhibited by insulin activation (Singh et al., 2018).

In gut epithelial cells, endoreplication is regulated by both Tor-dependent InR signaling and Tor-independent EGFR/MAPK signaling (Xiang et al., 2017). Under starvation conditions, Hh is released from enterocytes (Rodenfels et al., 2014), activating several genes controlling endoreplication (Fuß et al., 2001).

Using larvae reared in nutrient-restricted medium to inhibit InR signaling, we identified a potential role of Tor-independent mechanisms in promoting the muscular endocycle. Interestingly, a homolog of the transcription factor *Ci* (Gli) of the Hh pathway was identified in human glioblastoma cells as a positive regulator of Insulin Receptor Substrate (IRS, homolog of *chico*), which mediates cellular responses to insulin and insulin-like growth factor 1. The inactivation of *Gli* inhibits the MAPK-dependent insulin signaling pathway (Hsieh et al., 2010; Machado-Neto et al., 2018).

Furthermore, mild muscle mitochondrial distress may be associated with increased lifespan via the potential repression of systemic insulin signaling as suggested by previous studies (Owusu-Ansah et al., 2013). Therefore, the increased lifespan and the muscular phenotype observed here in *Ptr*^*23c*^ larvae suggest that they could have a MAPK-dependent pathway enhanced or some inhibition released.

Potential interactions between the InR/FOXO pathway, Hh signaling, and insulin-IGF pathways, in line with existing literature (Hwangbo et al., 2004; Singh et al., 2018) might underlie the phenotype we observed in the *Ptr*^*23c*^ mutant. However, the intricacies of these interactions require further exploration.

## Conclusions

Our study of the mutant phenotype *Ptr*^*23c*^, a null mutation in *Ptr*, indicated that this gene has a multifaceted role throughout the entire life cycle. The high mortality during embryonic, larval and pupal stages, the deficient locomotion of the larvae and the unexpected extension of adult lifespan offer a complex picture that requires further exploration. Our findings also open avenues for investigating the broader genetic networks governing these intricate processes.

## Acknowledgments

The authors would like to thank Prof. Em. Rafael Cantera for his continued support and critical reading of the manuscript; Dr. María José Ferreiro for fruitful discussions and expert advice; Dr. Rosa Barrio for kindly providing the *w;Df(2L)32FP-5/CyO*,*twist-Gal4*,*UAS-GFP* stock, and Dr. Carmen Bolatto for kindly providing the *Ptr*^*23c*^*/CyO, ftz-lacZ* stock. This work was partially funded by a CSIC grant conferred to CP (CSIC Iniciación a la Investigación, 2017, ID216), funds from PEDECIBA (Programa de Desarrollo de las Ciencias Básicas) and funds granted to the Departamento de Biología del Neurodesarrollo (IIBCE, Montevideo). Support from Agencia Nacional de Investigación e Innovación (ANII) to DP through Sistema Nacional de Investigadores (SNI) is gratefully acknowledged.

